# Methylation and Transcriptomic Profiling Reveals Short Term and Long Term Regulatory Responses in Polarized Macrophages

**DOI:** 10.1101/2024.06.18.599278

**Authors:** Giorgia Migliaccio, Jack Morikka, Giusy del Giudice, Maaret Vaani, Lena Möbus, Angela Serra, Antonio Federico, Dario Greco

**Author notes:** Corresponding author, FHAIVE, Faculty of Medicine and Health Technology, University of Helsinki. Arvo Ylpönkatu 34, 33100 Tampere.

## Abstract

Macrophage plasticity allows the adoption of distinct functional states in response to environmental cues. While unique transcriptomic profiles define these states, focusing solely on transcription neglects potential long-term effects. The investigation of epigenetic changes can be used to understand how temporary stimuli can result in lasting effects. Moreover, epigenetic alterations play an important role in the pathophysiology of macrophages, including phenomena related to the trained innate immunity, which allow faster and more efficient inflammatory responses upon subsequent encounters with the same pathogen. In this study, we used a multi-omics approach to elucidate the interplay between gene expression and DNA-methylation, unravelling the long-term effects of diverse polarizing environments on macrophage activity. We identified a common core set of genes that are differentially methylated regardless of exposure suggesting a potential mechanism for rapid adaptation to various stimuli. These conserved epigenetic modifications might represent a fundamental state that allows for flexible responses to various environmental cues. Functional analysis revealed that processes requiring rapid responses displayed transcriptomic regulation, whereas functions critical for long-term adaptations exhibited co-regulation at both transcriptomic and epigenetic levels. Our study unveils a novel set of genes critically linked to the long-term effects of macrophage polarization. This discovery underscores the potential of epigenetics in elucidating how macrophages establish long-term memory and influence health outcomes.

**Highlights:**

- Environmental signals trigger gene changes in macrophages, leaving a long-lasting epigenetic reprogramming
- Epigenetic changes and metabolic shifts in polarized macrophages suggest training mechanisms
- Common gene set epigenetically altered across different cues, suggest common adaptation to various stimuli

**Graphical Abstract:** 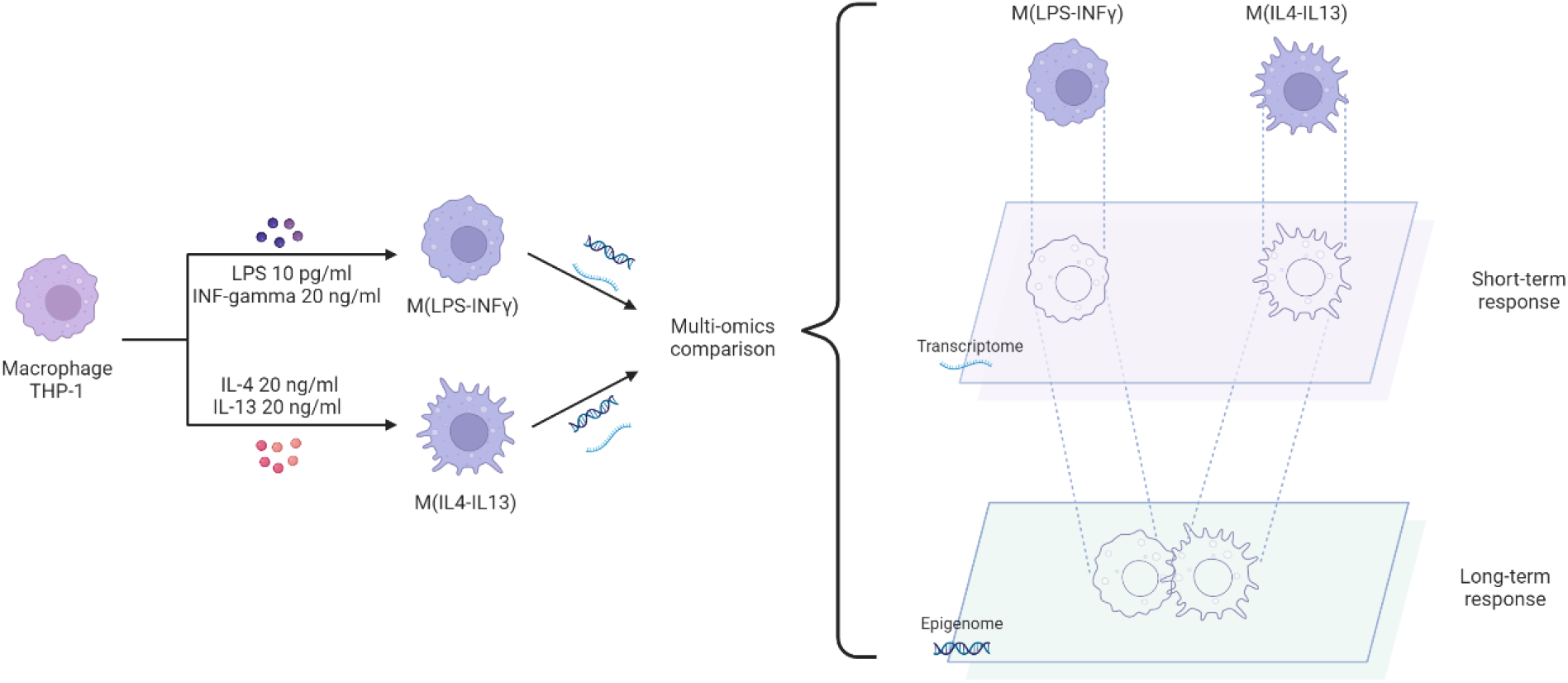

## 1. Introduction

Macrophages are a versatile cellular component of the immune system and are one of the first lines of defense of the body against pathogens or tissue damage (1). The plasticity of macrophages allows them to adopt distinct functional states in response to microenvironmental cues (2). The traditional classification of classically activated or pro-inflammatory (PI), induced by bacterial lipopolysaccharide (LPS) and/or interferon-gamma (IFNγ), and alternatively activated macrophages or anti-inflammatory (AI), that can be triggered by, for example, interleukin-4 (IL-4), represents the extremes of this functional spectrum (3). This spectrum of activation allows macrophages to exhibit functional diversity, playing crucial roles in pathophysiological processes and responses to foreign materials (4,5). Macrophage plasticity is tightly regulated by transcriptional reprogramming where macrophage phenotypes can be defined by distinct gene expression profiles (6–8). Epigenetic shifts are increasingly recognized as directing the transcriptional differences observed between macrophages in polarized states (9–12). Epigenetic modifications such as DNA methylation bridge the gap between transient stimuli and long-lasting cellular responses, even after the initial stimulus has subsided (11,13–15). Of all types of epigenetic modification, DNA methylation is the most stable and persistent (16,17). Epigenetic, transcriptional, and metabolic reprogramming is responsible for the protective effects of trained immunity (18). Trained immunity describes the long-term functional reprogramming of innate immune cells in response to environmental cues, which leads to an altered response to further challenges (19–21). The demanding energy requirements for an immune response to foreign insults necessitate the coordinated regulation of multiple metabolic pathways to act as sources of essential metabolites that are required for epigenetic alterations, serving as cofactors for chromatin-modifying enzymes (22,23).

Integrating transcriptomic and DNA methylation data allows to elucidate how macrophage polarization leads to long-term reprogramming for subsequent challenges. This combined analysis can shed light on whether biological or synthetic insults can induce long-lasting epigenetic imprinting, while also revealing co-regulatory pathways and genes critical for the trained macrophage response (24,25). While traditional toxicology focused on observable phenotypic effects, the field is increasingly embracing toxicogenomics to elucidate the mechanisms of action of chemicals. THP-1 macrophages, a well-established cell model for innate immunity testing in traditional toxicology (e.g. https://one.oecd.org/document/env/cbc/mono(2022)16/en/pdf), are now being explored for their utility in toxicogenomic safety assessments. Previous work, for example, has integrated DNA methylation changes and gene expression changes to investigate the effect of carbon nanotube exposure both in vitro and in vivo, discovering genes and pathways regulated at both levels linked to the outcome of lung fibrosis (15,26). Other studies focusing on polarized macrophages have also explored different types of epigenetic control such as posttranslational histone modifications and chromatin accessibility (6,10,11) and whilst there have been explorations of general DNA methylation dynamics in immune cells (27,28), studies looking at differences in methylation patterns between distinctly polarized macrophage states are still missing.

Macrophages respond to micro-environmental changes mediated by cytokines and other molecules secreted by other cells as well as pathogens. This dynamic interplay between macrophages and their environment creates a complex communication network that ultimately shapes their effector function (2). Here we employed a multi-omics approach, integrating DNA methylation profiling with transcriptomic analysis of polarized macrophages. Macrophages were polarized into distinct phenotypes using LPS and IFNγ established PI inducers LPS and IFNγ (29) and interleukin-4 (IL-4) and interleukin-13 (IL-13), known to promote an AI phenotype (M(IL4-IL13))(4). We identified and functionally characterized the set of genes that are differentially expressed or epigenetically modified in these M(LPS-IFNγ) and M(IL4-IL13) macrophages. LPS and IFNγ triggered a significantly higher number of differentially expressed genes (DEGs) compared to IL4-IL13. At the epigenetic level, however, both stimuli predominantly affected the same genes, suggesting a common core set of genes that are differentially methylated regardless of the environmental cue at each end of the polarization spectrum. We also identified a smaller subset of genes displaying environment-specific epigenetic regulation, highlighting a potential for distinct regulatory mechanisms at the level of DNA methylation changes. At the functional level, immune and metabolic processes were regulated epigenetically in both polarized states, imprinting the need for potential long-term shifts in immune and metabolic activity. At the transcriptomic level, the same metabolic processes were regulated in the PI phenotype but not in the AI phenotype, suggesting fast metabolic shifts are necessary in response to acute inflammatory insults, but in an AI state the macrophages are instead primed for potential future metabolic activity changes. Building upon the established link between metabolic rewiring and epigenetic reprogramming in trained immunity, this study offers a valuable framework to investigate the mechanisms underlying training in polarized macrophages. In modern toxicology, multi-omic strategies are gaining traction for their ability to provide a more comprehensive picture of cellular responses. We applied this strategy on THP-1 macrophages, a well-established *in vitro* model system, to identify differentially regulated processes at the transcriptomic and DNA methylation levels according to their polarization state. This approach elucidates both short-term and long-term regulatory strategies, informing our understanding of macrophage population responses across the polarization spectrum in pathophysiological conditions and following chemical exposures.

## 2. Materials and Methods

### 2.1 THP-1 cell culture

THP-1 cells (ATCC TIB-202, USA) were cultured in RPMI-1640 (Gibco, USA) supplemented with 10% FBS (Gibco, USA) (culture media). Cells were cultured in 75 cm^2^ flasks at a density < 1 × 10^6^ cells/ml. Cells were differentiated in 12 well plates, with 500,000 cells/ well (127,000 cells/cm^2^), in the culture media supplemented with 30.9 ng/ml of phorbol 12-myristate 13-acetate (PMA) (Sigma-Aldrich, USA) for 48h. After PMA differentiation, cells are exposed to two different cytokines cocktails and fresh media for 24h, 48h, and 72h. In particular, to simulate a PI environment the cells are exposed to LPS (Merck) 10 pg/ml and IFNγ (Sigma-Aldrich, USA) 20 ng/ml; to induce an AI phenotype instead, the cells are exposed to IL-13 (Sigma-Aldrich, USA) 20 ng/ml and IL-4 20 (Sigma-Aldrich, USA) ng/ml.

### 2.2 Collection of RNA

Cells were lysed on the plate and the RNeasy mini kit (Qiagen, Germany) was used as per the vendor’s instructions. The quality of the RNA samples was verified using Bioanalyzer 2100 (Agilent, USA), with RNA 6000 Nano Kit (Agilent, USA) following the vendor’s instructions. The RIN values for each sample exceeded 9.0.

### 2.3 RNA-sequencing

Quantity and quality of the RNA samples were assessed with quality checks as follows: preliminary quality control was performed on 1% agarose gel electrophoresis to test RNA degradation and potential contamination, sample purity and preliminary quantitation and RNA integrity were measured using Bioanalyzer 2100 (Agilent Technologies, USA). For library preparation, the Novogene NGS RNA Library Prep Set (PT042) was used. The mRNA present in the total RNA sample was isolated with magnetic beads of oligos d(T)25 using polyA-tailed mRNA enrichment. Subsequently, mRNA was randomly fragmented and cDNA synthesis using random hexamers and reverse transcription was performed. Once first chain synthesis was finished, the second chain was synthesised with the addition of an Illumina buffer (non-directional library preparation). Together with the presence of dNTPs, RNase H and polymerase I from E. Coli, the second chain was obtained by Nick translation. The resulting products then underwent purification, end-repair, A-tailing and adapter ligation. Fragments of the appropriate size were enriched by PCR, where indexed P5 and P7 primers (Illumina) were introduced, and final products were purified. The library was checked with Qubit 2.0 and real-time PCR for quantification and bioanalyzer Agilent 2100 for size distribution detection. Quantified libraries were pooled and sequenced on the Illumina Novaseq X platform, according to effective library concentrations and data amounts using the paired-end 150 strategy (PE150).

### 2.4 RNA-sequencing data analysis: data collection and pre-processing

The “.fastq” format raw files were used for the quality assessment of all RNA-Seq datasets, and it was conducted using FastQC v0.11.7. Trimming of reads, including removal of low-quality ends and adapters, was performed using cutadapt v4.4_dev with Python 3.8.10 with default parameters. Subsequently, the trimmed and adapter-clipped raw reads underwent a second round of quality checks using FastQC v0.11.7. The RNA sequencing reads were aligned to the human reference genome assembly GRCh38 using the HISAT2 algorithm and its corresponding genome indexes. Transcript abundance was determined using the featurecounts function from the Rsubread v1.34.4 R package with Rstudio version 1.1.453, using Ensembl annotation version 105 (https://www.ensembl.org). Non-expressed and lowly expressed genes were filtered out by applying a proportion test, as implemented in the NOISeq Bioconductor package.

### 2.5 RNA-sequencing: Differential expression analysis

The DESeq2 v1.24.0 R package was employed for normalization and differential expression analysis (30) (the data are deposited in Zenodo with the following doi 10.5281/zenodo.11473632). Specifically, we conducted a comparison for differential gene expression at each time point between the treatments and the control (e.g. M(LPS-IFNγ)_24h vs control_24h, M(LPS-IFNγ)_48h vs control_48h, etc.), as well as for the differential expression of genes among control groups over time (e.g. control_48h vs control_24h). The analysis was performed by using the cytokines cocktail and time of exposure together as variables of interest. Differentially expressed genes (DEGs) are determined by comparing samples from the treatment group to untreated samples the genes are then filtered for adjusted p-value FDR<0.01 and absolute log2FC > 0.58. As our cell population comprises monocyte-derived macrophages, we took precautions to exclude any variations in gene expression linked to macrophage differentiation. Our focus remained solely on assessing the impact of the stimulation cocktail and we wanted to mitigate the potential impact of macrophage differentiation on our analysis. Consequently, genes displaying differential expression over time among the control groups (e.g. control_48h vs control_24h, control_72h vs control_24h) were omitted from the analysis.

### 2.6 DNA-methylation: data collection and pre-processing

Genomic DNA was extracted using a DNeasy Blood and Tissue Kit (Qiagen). The quality and the concentration of the DNA samples were verified using Nanodrop2000. Genomic DNA was treated with sodium bisulfite using EZ-96 DNA methylation kit (Zymo Research, cat no.: D5004), following the manufacturer’s standard protocol. Assessment of levels of DNA methylation of known CpG regions and promoters across the genome was done with Infinium MethylationEPIC v2.0 Kit (Illumina, Inc.) and Illumina iScan. In brief, following bisulfite conversion, approximately 500 ng of the bisulfite-converted DNA per sample was used for methylation analysis. The initial quality control and identification of signal intensities for each probe were performed with Illumina GenomeStudio Software.

Methylation data were analyzed in R following the workflow previously described by Maksimovic *et al.* (31). Raw methylation files were uploaded together with the metadata and microarray annotation file. Illumina methylation data is usually obtained in the form of Intensity Data (IDAT) files, a proprietary format that is generated by the scanner and stores summary intensities for each probe on the array. The raw intensity signals were read into R from the IDAT files using the “read.metharray.exp” function of the minfi package v1.46 (32). The function creates an RGChannelSet object that contains all the raw intensity data, from both the red and green color channels, for each of the samples. The quality of the dataset was evaluated first by calculating the detection p-value for every CpG in every sample and then generating a quality control report using the “qcReport” function of the minfi package (32). CpG probes were filtered by removing probes with a detection p-value higher than 0.05 in any sample (the data are deposited in Zenodo with the following doi 10.5281/zenodo.11473632). Further filtering was applied to remove probes for CpGs located on the sex chromosomes or those containing single nucleotide polymorphisms. To minimize the unwanted variation between samples, data were normalized using the “preprocessQuantile” function of the minfi package. Once the data had been filtered and normalized, the M-values and beta values were calculated. M-values have better statistical properties and are thus better for use in statistical analysis of methylation data whilst beta values are easy to interpret and are thus better for displaying data. M-values were obtained using the “getM” function from the minfi package (32), while the beta values were obtained using the function “getBeta” from the same package. Probe wise differential methylation analysis was performed using the beta values between each treatment and its control for each time point. The analysis was performed on the matrix of M-values using the limma package (33), obtaining moderated t-statistics and associated p-values for each CpG site, which has been further corrected with the FDR method. Given the genome-wide nature of the analysis and the inherent uncertainty associated with interpreting beta values, a stringent p-value threshold of 10^-8^ was employed to have a more robust statistical significance (34).

### 2.7 DNA-methylation: probe annotation

At the epigenetic level, probes showing differential methylation (FDR<10^-8^) between the control group over time were excluded. After pinpointing these probes, the GREAT software (35) was used for CpG annotation to genes (FDR<0.5). Specifically, each gene was allocated a regulatory domain extending to the midpoint between its transcription start site (TSS) and the nearest gene’s TSS, limited to 1000kb (**Supplementary Figure 1**). To ensure robust annotation, only genes with a minimum of 3 significant CpGs annotated were considered as differentially methylated genes (DMGs).

### 2.8 Functional annotation analysis

Pathway enrichments were performed using the “gost” function of the gProfiler package (36). Lists of official genes were offered as input after being converted to gene symbols. Gene ontology (GO) terms were enriched using all known genes as the statistical domain scope of the analysis. GO terms were considered significantly enriched with an FDR-adjusted p-value <0.05 (**Supplementary Table 1-2**). When comparing the functional annotation of the two phenotypes (e.g. M(LPS-IFNγ) and M(IL4-IL13)), the most significant terms are categorized into four groups: (a) unique to M(LPS-IFNγ), (b) unique to M(IL4-IL13), (c) shared and most significant in M(LPS-IFNγ), and (d) shared and most significant in M(IL4-IL13). This categorization will allow the identification of functionally distinct terms associated with each phenotype.

## 3. Results and discussion

### 3.1 Differential methylation patterns in polarized macrophages are numerically similar despite transcriptomic divergence

To evaluate quantitative perturbations in the transcriptome of macrophages treated with either LPS and IFNγ (M(LPS-IFNγ)), or IL4 and IL13 (M(IL4-IL13)) and and to assess whether this polarization resulted in distinct gene expression profiles, a differential expression analysis was performed. We identified 5,753, 6,209, and 5,021 DEGs following exposure to LPS and IFNγ, compared to untreated samples, at 24h, 48h, and 72h, respectively (**Figure 1A** in red). Similarly, macrophages treated with IL4 and IL13 exhibited altered gene expression patterns. The number of DEGs identified at 24h, 48h, and 72h were 1,527, 2,309, and 1,916, respectively (**Figure 1A** in green) (**Supplementary Table 3**). M(LPS-IFNγ) macrophages exhibit over twice as many DEGs compared to M(IL4-IL13) at all time points, revealing a divergence in gene expression between PI and AI macrophages consistent with previous findings (37–39). As LPS and IFNγ simulate the acute immune response to infection, while IL4 and IL13 are expressed during resolution of the acute phase, it suggests that the transcriptomic regulation of fewer genes is required for maintaining the AI state compared to the acute PI state. When examining the differential methylation results, the macrophages polarized in an environment rich in LPS-IFNγ do not exhibit any statistically significant DMGs at 24h but show 3,595 genes at 48h and 5,247 genes at 72h (**Figure 1B**). Similarly, the M(IL4-IL13) demonstrates differential methylation in 11 genes at 24h, and 2935 and 5241 genes at the later time points of 48h and 72h, respectively (**Figure 1B**) (**Supplementary Table 3**).

**Figure 1:**
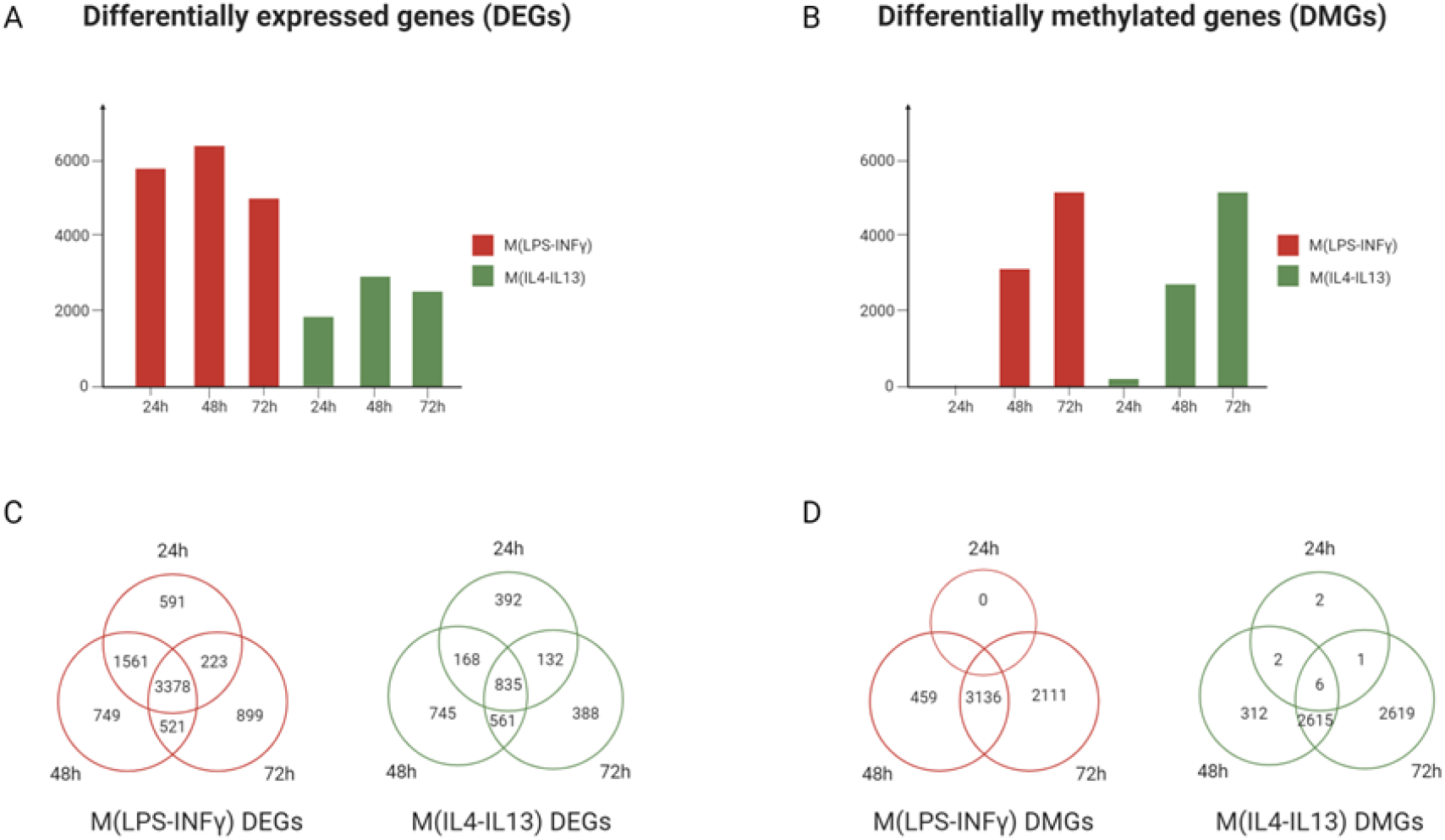
(A) Number of differentially expressed genes (DEGs) and (B) differentially methylated genes (DMGs) at time points 24H, 48H and 72H of pro-inflammatory stimulation (in red) and anti-inflammatory stimulation (in green). (C-D) Venn diagrams of DEGs and DMGs in time in the pro-inflammatory phenotype (in red) and anti-inflammatory phenotype (in green).

While the number of DMGs remains relatively similar between the two phenotypes, there is a significant divergence in the number of DEGs between the two conditions at each time point (**Figure 1A, 2B**). Analyzing the temporal consistency of differential expression (**Figure 1C**), some genes exhibit sustained changes across all three time points (24h, 48h, and 72h), in particular, 3,378 genes within the M(LPS-IFNγ) phenotype and 835 within M(IL4-IL13), that indicates a persistent variation in gene expression induced by the stimuli. Additionally, in both phenotypes, at the methylation level, a progressive increase in the number of DMGs over time is observed (**Figure 1B**), and most of the genes that are differentially methylated during the early time points are maintained for up to 72h (**Figure 1D**). Unlike the rapid fluctuations observed in gene expression, DNA methylation patterns are far more stable over short experimental time periods and take longer to shift in response to an exposure (40–43).

The influence of LPS and IFNγ on the transcriptome seems considerably stronger compared to IL4 and IL13, as indicated by a significantly higher number of dysregulated genes. However, this difference is not reflected at the epigenetic level, where the number of DMGs under the two conditions remains relatively similar. To understand the functional implications of the observed differences in gene regulation, functional annotation of the DEGs and DMGs was performed.

### 3.2 Functional annotation of transcriptome shows polarized phenotypes

To gain a deeper understanding of the biological processes affected by our polarization protocol, enrichment analysis has been performed to functionally annotate the DEGs. This analysis identified enriched GO terms, providing insights into potential functional patterns associated with polarized macrophages.

Enrichment analysis of the DEGs at 24h, 48h, and 72h in response to both cytokine-rich environments (i.e. LPS-IFNγ and IL4-IL13), revealed that the top 5 significant enriched terms remain consistent across all three time points. This suggests sustained functional changes across 72h of continuous exposure (**Supplementary Figure 1**). Therefore, we focused on the combined set of DEGs from all three time points. A total of 7,923 genes exhibited differential expression at the union of all three time points in macrophages stimulated with LPS and IFNγ compared to unexposed macrophages, whereas 3,222 DEGs were identified in macrophages exposed to IL4 and IL13. The comparison of DEGs between the two phenotypes revealed 2,720 genes commonly dysregulated in both M(LPS-IFNγ) and M(IL4-IL13), suggesting a subset of genes to be responsive to both stimuli (**Figure 2A**). To understand the directionality of these changes, we calculated the median fold change across all three time points in both exposures. This analysis identified 2,075 genes exhibiting concordant regulation (changing in the same direction) upon exposure to both stimuli, while 645 displayed discordant regulation between M(LPS-IFNγ) and M(IL4-IL13) (**Supplementary Table 4**). These 645 discordantly regulated genes were enriched in functional categories associated with cytokine signaling and inflammatory pathways (**Supplementary Table 4**). This enrichment aligns with the established roles of LPS-IFNγ and IL4-IL13 in driving distinct macrophage phenotypes with contrasting cytokine profiles and inflammatory responses. Additionally, 5,203 and 502 genes displayed unique dysregulation in M(LPS-IFNγ) and M(IL4-IL13) (**Figure 2A**), respectively, highlighting environment-specific transcriptional signatures. We conducted a gene ontology enrichment analysis, revealing distinct terms associated with each phenotype (***Figure 2*B**).

**Figure 2:**
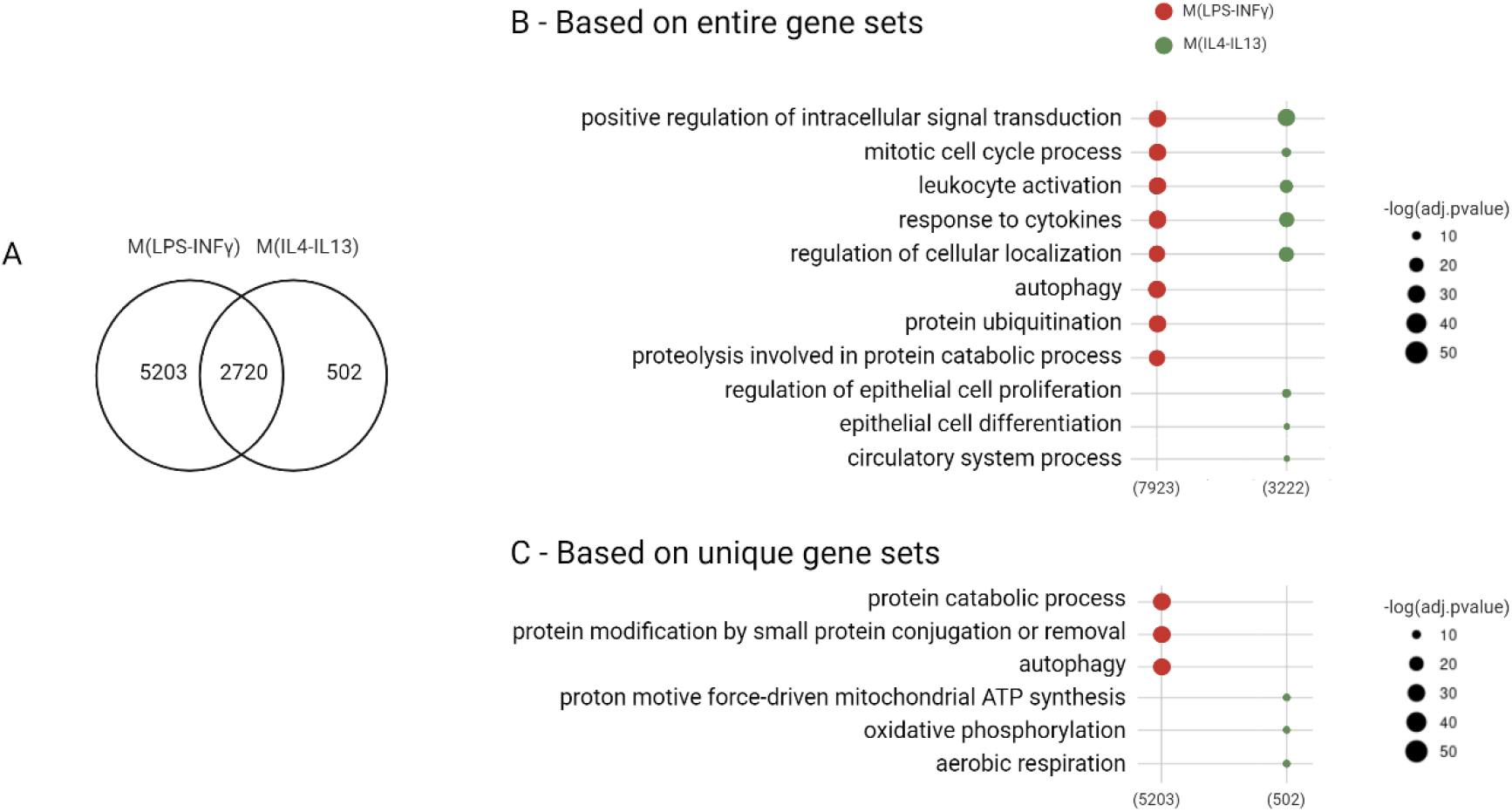
(A) Number of DEGs (FDR<0.01; abs(log2FC)>0.58) between macrophages exposed to LPS-IFNγ and IL4-IL13. (B) Gene ontology biological processes enrichment (FDR<0.05; terms with number of genes 5<N<1,000) is performed with the entire set of genes associated with each phenotype; the plotted terms are the three most significant ontologies in the category of shared and unique pathways. For the common pathways, the most significant three for each phenotype have been selected. (C) The enrichment analysis is performed within the gene set exclusive to each phenotype, not shared between the two (i.e. 5,203 and 502 respectively for M(LPS-IFNγ) and M(IL4-IL13)); the plotted terms are the three most significant ontologies in the category of unique pathways.

Consistent with the use of cytokines for polarization, functional enrichment analysis revealed immune pathway activation (e.g., “responses to cytokines”, “leukocyte activation”) in both M(LPS-IFNγ) and M(IL4-IL13) phenotypes (**Figure 2B**). Notably, M(LPS-IFNγ) uniquely displayed enrichment of autophagy, protein ubiquitination, and proteolysis pathways, suggesting enhanced antigen presentation and a PI state, potentially reflecting their role in combatting pathogens (44–46). In contrast, M(IL4-IL13) macrophages displayed unique enrichment of pathways associated with cell proliferation and differentiation (**Figure 2B**). This suggests a functional role in tissue repair and remodeling, potentially through the release of growth factors like TGFβ1 and platelet-derived growth factor (PDGF), which promote epithelial cell proliferation and fibroblast activation (47–49). Recognizing a substantial overlap in the DEGs between the LPS-IFNγ and IL4-IL13 stimulated phenotypes (**Figure 2A**), a further enrichment analysis focused on genes exclusive to each phenotype (i.e., 5,203 for M(LPS-IFNγ) and 502 for M(IL4-IL13)) was conducted to mitigate the bias introduced by shared gene counts. The functional annotation genes revealed distinct metabolic profiles associated with macrophage phenotypes ***Figure 2*C**). LPS-IFNγ treatment enriched terms related to macromolecule and protein breakdown, potentially reflecting autophagic processes in degrading foreign material (50,51). Conversely, the AI phenotype (M(IL4-IL13)) displayed enrichment of oxidative phosphorylation terms, suggesting a metabolic shift. This aligns with reports of M(LPS-IFNγ) favoring glycolysis and M(IL4-IL13) relying more on oxidative phosphorylation, highlighting the metabolic reprogramming that accompanies macrophage polarization (52,53).

Our transcriptomic analysis indicates that the macrophages successfully polarized in response to environmental-specific stimuli. The DEGs identified in the LPS-IFNγ and IL4-IL13 treated groups displayed functional annotations consistent with the PI and AI phenotypes, respectively. The M(LPS-IFNγ) group shows enrichment of terms related to inflammation, autophagy, and immune cell activation, characteristic of the PI phenotype. Conversely, the M(IL4-IL13) group displays enrichment of terms associated with oxidative phosphorylation metabolism and tissue repair pathways, aligning with the AI phenotype.

### 3.3 Polarized macrophages show common and distinct epigenetic regulation

Through the examination of epigenetic modification derived from polarized macrophages, our goal is to discover the modifications occurring in DNA methylation levels when monocyte-derived macrophages are subjected to environmental cues mimicking the PI and AI conditions. Deciphering the unique epigenetic signatures associated with polarized macrophage states offers valuable insights into the long-term impact on immune function and their memory-like capabilities. Epigenetic regulation plays a significant role in trained immunity, a process where innate immune cells, like macrophages, exhibit memory that alters response to future encounters (54–57). While previous research explored epigenetic modification by investigating changes in the activity of molecules like DNA methyltransferases and histone modifiers (10,12,58), our study takes a novel approach by directly identifying sets of common and unique DMGs associated with polarized macrophages.

A progressive increase in DMGs is observed over time both in M(LPS-IFNγ) and M(IL4-IL13) (**Figure 1B**). Recognizing the slower dynamics of the DNA methylation process compared to gene expression fluctuations (17,40,42), genes that did not display persistent methylation status were filtered out (i.e. genes showing methylation status change at 24h and 48h that are not seen at 72h, were omitted as false positives). This approach yielded 4,856 and 4,899 DMG in the cells stimulated with LPS-IFNγ and IL4-IL13, respectively. In total, there are 4,419 DMGs shared between the two phenotypes, 437 uniquely differentially methylated in M(LPS-IFNγ) and 480 in M(IL4-IL13) (**Figure 1A**). Our investigation of DMGs between phenotypes did not explore the directionality (e.i. hypomethylation and hypermethylation) of these genes. This decision reflects the ongoing challenge of elucidating the precise impact of methylation or demethylation at a single CpG on overall gene expression (59). The presence of multiple CpG sites within a single gene further complicates this interpretation.

Our results suggest that, despite the polarization phenotype, macrophages retain a core set of DNA methylation changes, suggesting a potential mechanism for rapid adaptation to various stimuli. These conserved epigenetic modifications might represent a fundamental state allowing flexible responses to various environmental cues. These findings show the presence of a dual regulatory strategy: a bigger common core set of DMGs and a second regulation of DMGs responding to the specific initial stimulus.

Functional analysis of DNA methylation changes (i.e. 4,856 genes in M(LPS-IFNγ) and 4,899 in M(IL4-IL13)) revealed shared enrichment for DNA damage response and non-coding RNA (ncRNA) regulation (**Figure 3B**). The shared enrichment for DNA damage response pathways in both M(LPS-IFNγ) and M(IL4-IL13) phenotypes was unexpected. While LPS exposure is known to induce genotoxic stress via TLR activation (66, 67), this finding suggests that the M(IL4-IL13) phenotype also experiences some kind of genotoxic stress through an unidentified mechanism or that these cells are primed for a more robust DNA damage response upon future exposures. Furthermore, ncRNA regulation suggests epigenetic reprogramming, potentially underlying mechanisms of trained immunity. This aligns with existing evidence demonstrating ncRNA involvement in modulating immune responses and shaping long-lasting epigenetic modifications (60–63). Our analysis also revealed a small set of distinct functional terms associated with each macrophage phenotype (**Figure 3B, C**), suggesting an epigenetically imprinted response to environmental cues. Notably, M(LPS-IFNγ) exhibited enrichment for terms related to chromatin and histone modifications. This finding strengthens existing knowledge about the connection between metabolic pathways and epigenetic changes. For example, LPS stimulation is known to increase metabolites like cytosolic acetyl-CoA, which can facilitate histone acetylation at promoters and enhancers of LPS target genes such as IL6 and IL12B (64–67). This demonstrates how metabolic reprogramming induced by LPS exposure can directly influence chromatin structure and gene expression through epigenetic modifications. Conversely, the M(IL4-IL13) phenotype showed enrichment for “membrane organization” terms and lysosomal/lytic compartments (**Figure 3B, C**). While the specific link between this phenotype and metabolism requires further investigation, the dynamic nature of these compartments suggests the potential for metabolic regulation and digestion of pathogens (68–70). The observed epigenetic regulation of genes governing lysosomal membrane dynamics suggests a potential role for the AI environment in modulating macrophage metabolism, potentially influencing antigen processing during subsequent challenges.

**Figure 3:**
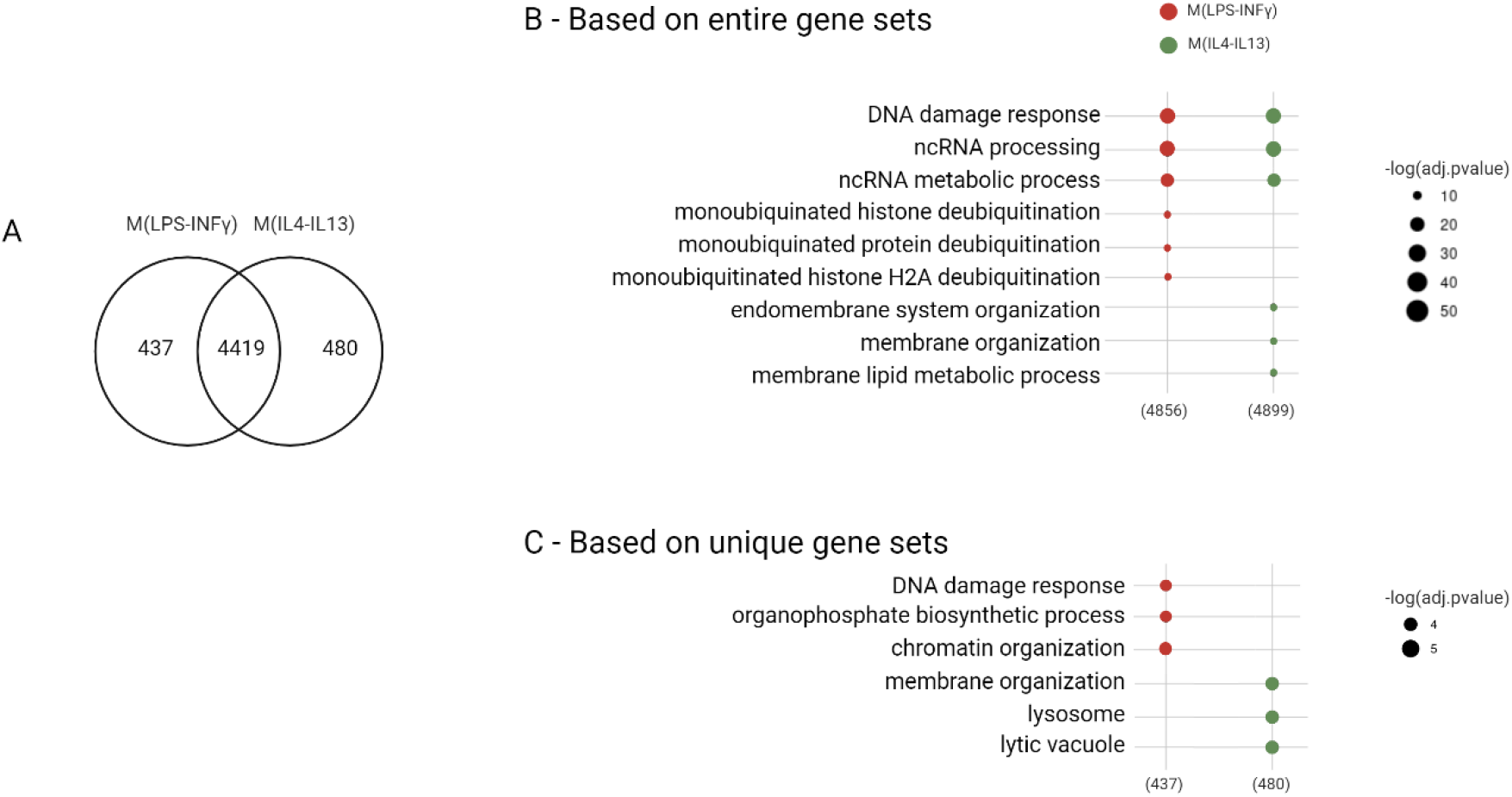
(A) Number of differentially methylated genes (DMG) (FDR<0.05) between macrophages exposed to pro-inflammatory stimuli M(LPS-IFNγ) and anti-inflammatory stimuli M(IL4-IL13). (B) Gene ontology biological processes enrichment (FDR<0.05; terms with the number of genes 5<N<1,000) is performed with the entire set of genes associated with each phenotype; the plotted terms are the three most significant ontologies in the category of shared and unique pathways. For the common terms, the most significant three for each phenotype have been selected. (C) The enrichment analysis is performed within the gene set exclusive to each phenotype not shared between the two (i.e. 437 and 408 respectively for M(LPS-IFNγ) and M(IL4-IL13)); the plotted terms are the three most significant ontologies in the category of unique pathways.

Our analysis of the epigenetic landscape in M(LPS-IFNγ) and M(IL4-IL13) macrophages revealed the presence of a dual regulatory strategy. A larger core set of DMGs is regulated regardless of the initial stimulus and likely regulates macrophage functions that are common across different polarization states. This shared core suggests epigenetic priming that is stimulus-independent. The second strategy of regulation exists through environmental-specific DMGs in the spectrum of PI and AI polarization. Their presence suggests a long-lasting epigenetic memory imprinted by the initial polarization signal that may extend beyond the immediate effects and potentially influence the long-term fate and function of the cells. By identifying these epigenetically regulated genes, this dataset provides valuable insights not only into macrophage polarization, but also into the process of macrophage training.

### 3.4 Integrated analysis of transcriptomic and epigenetic alteration unveils distinct regulatory mechanisms of macrophage activity in polarized states

After confirming effective polarization via transcriptome analysis and observing both commonalities and differences in the epigenetic layer between the two phenotypes, a multi-omics approach was employed to gain a comprehensive understanding of environment-induced alterations. Unlike transcriptional changes, which often follow an ‘impulse-like’ kinetic, epigenetic modifications can persist even after the initial stimulus fades (41,71). This enduring nature of epigenetic alterations makes them better long-term indicators of exposure-induced changes as compared to analyzing transcript levels in constant flux. However, transcriptomic changes are also indicative of immediate cellular needs and are required to direct the shift in DNA methylation changes. Therefore, we aimed to combine analysis of both layers of gene expression regulation to develop a contextualized view of the regulatory relationships between dynamic and immediate transcriptomic regulation and long-term epigenetic regulation post exposure in polarized states of macrophage activation.

Following the enrichment analysis of DEGs and DMGs in each phenotype, the ontologies were categorized into three groups: transcriptomic-specific, epigenetic-specific, and shared regulation (enriched in both transcriptomic and epigenetic datasets). From each group, the 10 most significant terms were selected and grouped based on broader biological processes (i.e. *cell death*, *signaling*, *cell-ECM interaction*, *developmental process*, *cell cycle*, *cellular response to stimulus*, *immune system process*, *metabolic process*, *translation and protein modifications*, *gene expression regulation*) (**Figure 4**). Analysis of the enriched terms revealed primarily transcriptomic regulation for categories like *cell death*, *signaling*, and *cell-ECM interaction*. This suggests that these processes rely on rapid gene expression changes, rather than long-term epigenetic modifications, to ensure precise control and adaptability. Apoptosis, for instance, requires quick activation and reversal based on cellular threats, favoring dynamic transcriptional responses over long-lasting epigenetic regulation (72). Similarly, cell-ECM interactions require swift adjustments in gene expression for cell mobility within a dynamic environment (73,74). Interestingly, “collagen fibril organization” within cell-ECM interaction showed evidence of epigenetic regulation in both phenotypes. This suggests a potential role for epigenetic priming in macrophage-fibroblast interaction during the resolution phase of inflammation, as macrophages encounter a collagen-rich matrix (75).

**Figure 4:**
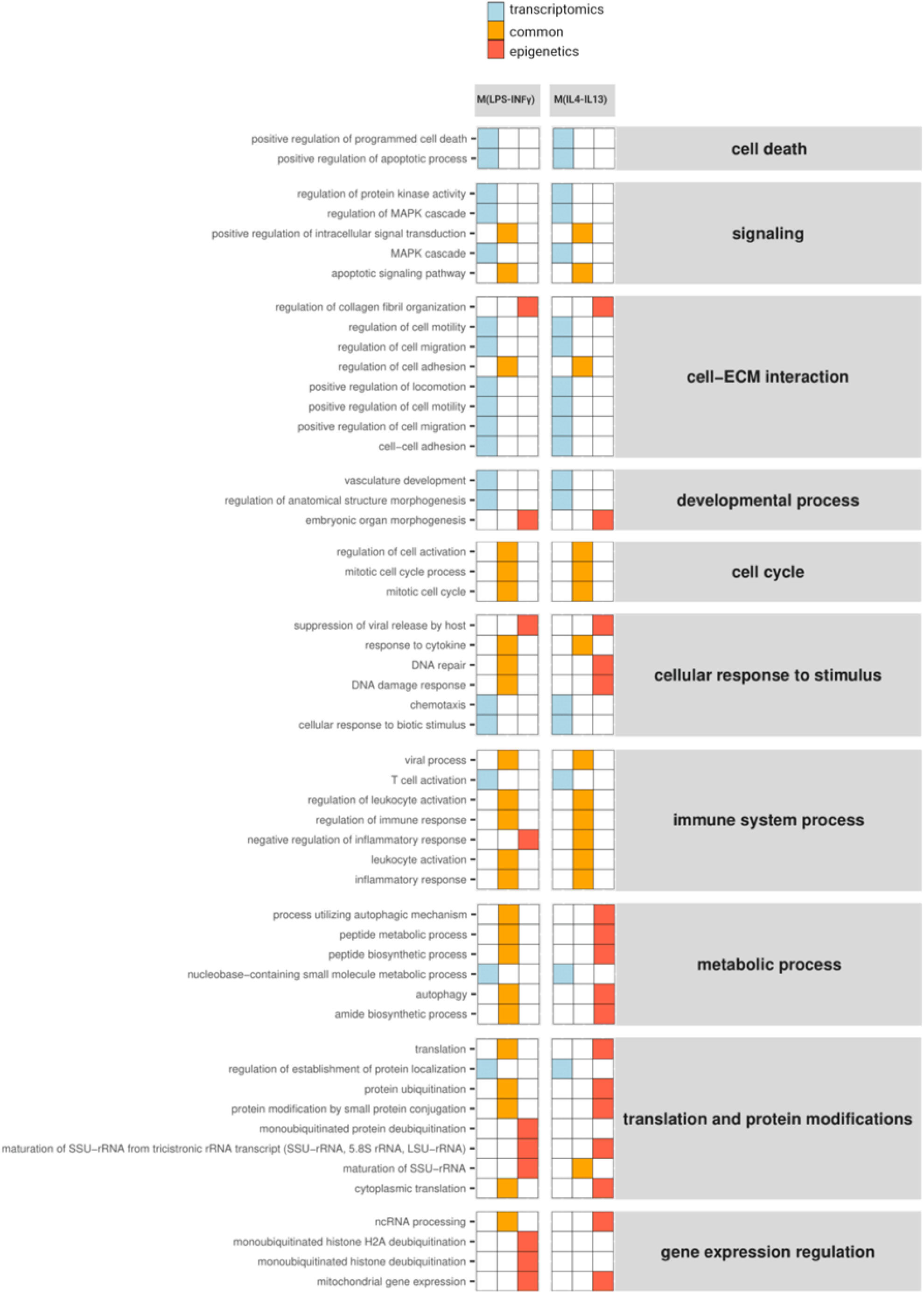
The integration of transcriptomic and epigenetic data by combined analysis of pathway enrichment of DEGs and DMGs. The top 10 most significant terms are presented for each phenotype, categorized into transcriptomics, epigenetics, or common terms. The p-value of common terms is evaluated with average FDR. Different colors (light blue, orange, red) distinguish these categories. The enriched pathways are grouped under broader biological processes (cell death, signaling, cell-ECM interaction, developmental process, cell cycle, cellular response to stimulus, immune system process, metabolic process, translation and protein modifications, gene expression regulation).

Immune system terms (**Figure 4**) displayed co-regulation at both transcriptomic and epigenetic levels in both phenotypes. This suggests rapid transcriptional changes for the initial immune response and epigenetic modifications for potential long-term memory. The observed epigenetic regulation across various immune-related terms (“inflammatory response”, “regulation of immune response”, etc.) indicates a potential form of cellular memory established following stimulation. This epigenetic reprogramming could prime macrophages for a more robust response upon future challenges, highlighting the integrated control of immune responses by rapid transcriptional modulation and long-term epigenetic memory.

Intriguingly, “DNA damage response” and “DNA repair” displayed contrasting regulation patterns (**Figure 4**). While M(LPS-IFNγ) exhibited co-regulation at both transcriptomic and epigenetic levels, M(IL4-IL13) showed solely epigenetic regulation. This aligns with the previously observed enrichment of DNA damage response pathways in both phenotypes (mentioned earlier). It suggests a potential priming mechanism for M(IL4-IL13) macrophages. The lack of immediate stress response (i.e., transcriptomic changes) in M(IL4-IL13) might be due to the absence of LPS-induced stress. However, epigenetic regulation of these genes could establish a “memory” for future stress challenges, potentially leading to a more efficient response (76). Protein production and metabolism terms (“process utilizing autophagic mechanism”, “peptide metabolic process”, etc.) exhibited contrasting regulation patterns (**Figure 4)**. M(LPS-IFNγ) exhibited co-regulation at both the transcriptional and epigenetic levels. This aligns with the established role of rapid transcriptional changes for a robust response during acute inflammation (77,78), further supported by the higher number of DEGs observed upon exposure (**Figure 1A**). In contrast, M(IL4-IL13) macrophages showed solely epigenetic regulation, suggesting a potential form of pre-adaptation. This epigenetic regulation could prime the transcriptomic machinery for a rapid metabolic shift if future environmental insults are encountered, potentially leading to a more efficient response.

## 4. Conclusion

This study investigated the long-term reprogramming induced by macrophage polarization, potentially establishing an epigenetic memory for subsequent challenges. This study supports the growing interest in toxicogenomics in understanding with multi-omics analysis, how various insults can have short-term effects and/or induce long-term epigenetic changes in immune cells. By employing multi-omics analysis, this strategy has unveiled co-regulatory pathways and key genes involved in the trained macrophage response, potentially impacting toxicogenomic safety assessments.

Transcriptomic analysis revealed distinct gene expression profiles between polarized macrophage states, reflecting their functional diversity. Interestingly, DNA methylation displayed a higher degree of similarity across these states. This finding suggests the existence of a core set of genes with stable epigenetic modifications, potentially independent of the initial environmental cue. By testing the identified core set across diverse stimuli, we can determine if these genes represent key regulators of macrophage plasticity and memory formation. While the precise role of individual CpG methylation in regulating gene activity remains an ongoing area of research, a deeper investigation into the methylation status of this core set and its influence on gene expression is necessary to fully understand the functional significance of these observations. Our multi-omics analysis suggests intriguing regulatory patterns across the two phenotypes. While some processes exhibited coordinated changes at both the transcriptome and epigenome levels, others displayed regulation solely at the epigenetic level. This observation prompts further investigation into the mechanisms driving these epigenetic modifications and the potential absence of a corresponding transcriptomic priming event. Furthermore, the multi-omics analysis showed that processes requiring rapid responses, such as motility and adhesion, exhibited regulation primarily at the transcriptomic level. Conversely, functions critical for long-term adaptations, like protein production, metabolism, and immune regulation, displayed co-regulation at both transcriptomic and epigenetic levels. This suggests that epigenetic modifications might play a pivotal role in establishing a long-term memory for these processes, particularly the observed metabolic shift and epigenetic alterations, considered hallmarks of trained immunity.

Overall, this study unveils the intricate interplay between transcriptomics and epigenetics in macrophage polarization. The identified core set of epigenetically modified genes presents exciting opportunities for novel therapeutic targets and enhances toxicogenomic safety assessments. Elucidating this spectrum across diverse stimuli, including other environmental cues, is crucial for advancing trained immunity research and developing targeted immunotherapies. This knowledge empowers future research to leverage epigenetics for therapeutic benefit and predict long-term health consequences of environmental exposures.

## Supporting information

Supplementary images and Tables

## Author contribution

G.M.: Conceptualization, Methodology, Validation, Formal analysis, Investigation, Writing - Original Draft, Visualization

J.M.: Methodology, Investigation, Writing - Original Draft, Writing - Review & Editing, Supervision

G.d.G.: Methodology, Formal analysis, Investigation, Writing - Review & Editing, Visualization

M.V.: Investigation, Writing - Review & Editing

L.M.: Writing - Review & Editing, Supervision

A.S.: Writing - Review & Editing, Visualization, Supervision

A.F.: Investigation, Writing - Review & Editing, Supervision

D.G.: Conceptualization, Methodology, Investigation, Resources, Writing - Review & Editing, Supervision, Project administration, Funding acquisition

## Data availability

Data are deposited in Zenodo with the following link https://doi.org/10.5281/zenodo.11473633.

## Declaration of Generative AI and AI-assisted technologies in the writing process

During the preparation of this manuscript, the authors utilized Google Gemini to improve the grammar and clarity of the text. After using this tool/service, the author(s) reviewed and edited the content as needed and take(s) full responsibility for the content of the publication.

