## Supplementary images and Tables for "Methylation and Transcriptomic Profiling Reveals Short Term and Long Term Regulatory Responses in Polarized Macrophages": Supplementary.docx

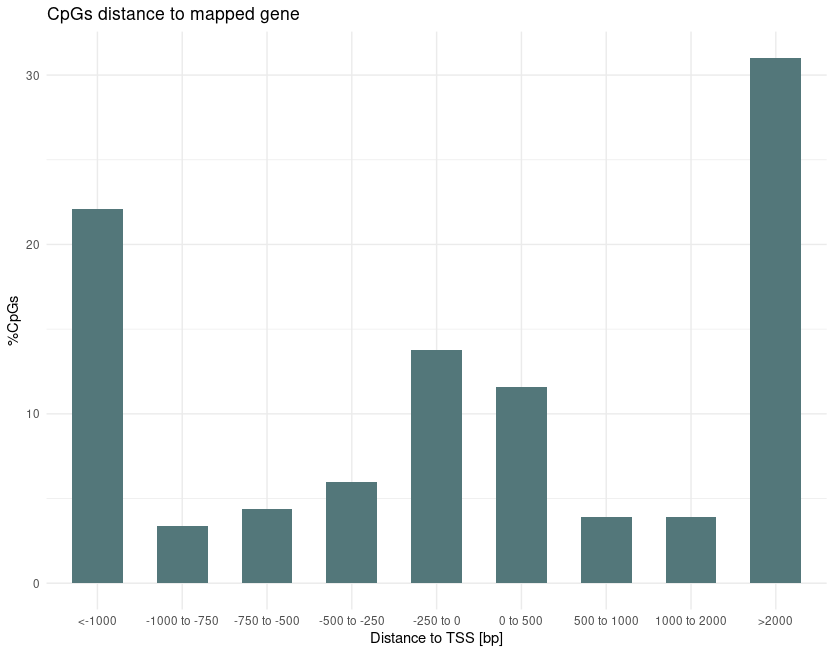

Supplementary Figure 1:	CpGs distance to TSS of mapped genes.

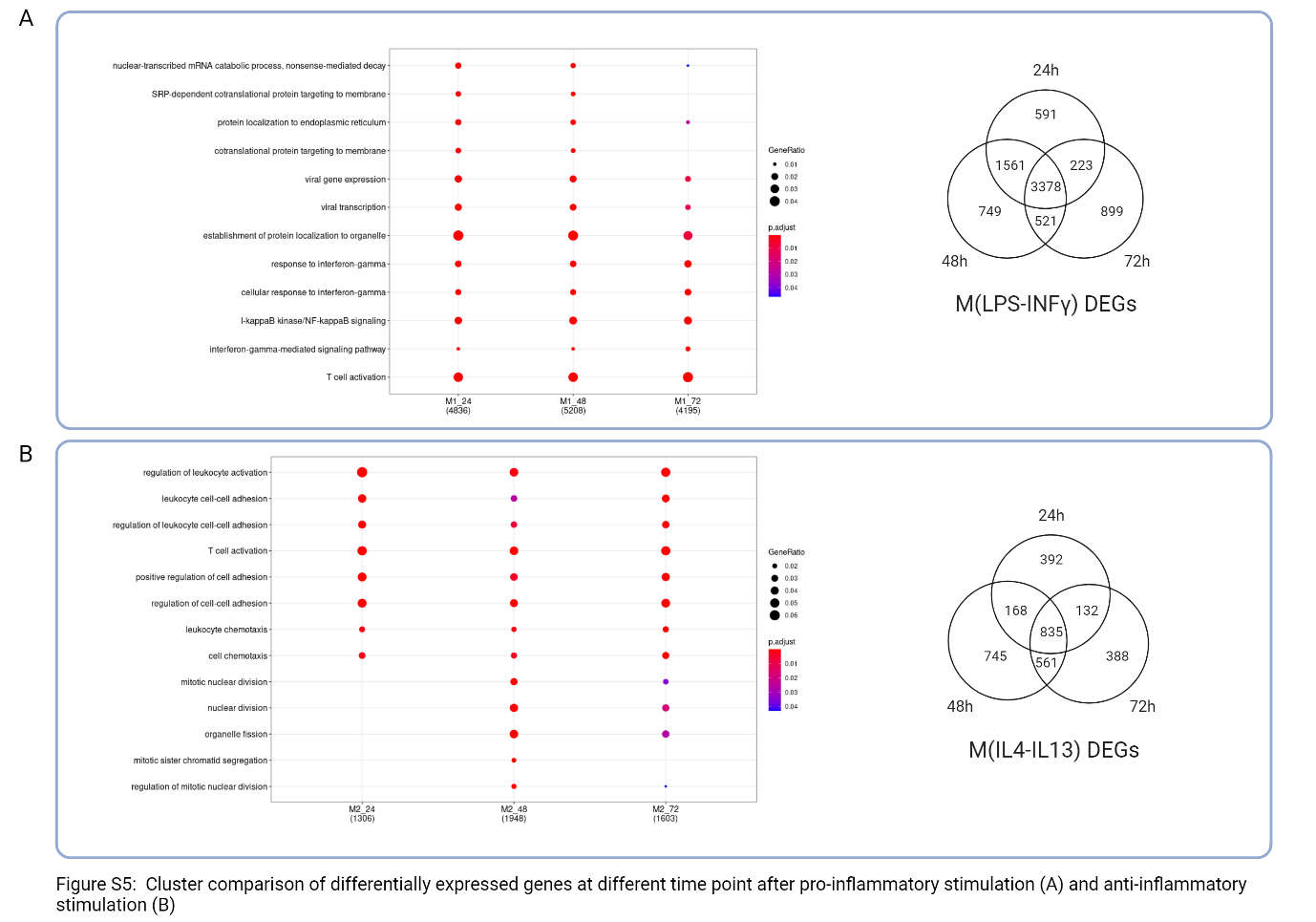

Supplementary Figure 2: Clusted comparison and Venn diagram of differentially expressed genes at 24h,48h and 72h after stimulation with LPS and IFNγ (A) and IL4-IL13 (B).

Supplementary Table 1: Enrichment analysis performed with gost function on the entire set of (Differentially Expressed Genes) DEGs and (Differentially Methylated Genes) DMGs associated with M(LPS-INFγ) and M(IL4-IL13).

Supplementary Table 2: Enrichment analysis performed with gost function on the set of DEGs and DMGs associated uniquely with M(LPS-INFγ) and M(IL4-IL13).

Supplementary Table 3: List of DEGs with FoldChange and pvalues from DESeq2 function.

Supplementary Table 4: Enrichment analysis performed with gost function on the common DEGs between the two phenotypes.
